# Oxytocin, Compulsion and Epilepsy: Insights from a Complex Behavioral and Neuronal Networks Association

**DOI:** 10.1101/638452

**Authors:** Simone S. Marroni, Victor R. Santos, Olagide W. Castro, Julian Tejada, Jessica Santos, Jose Antonio Cortes de Oliveira, Norberto Garcia-Cairasco

## Abstract

Previously we have demonstrated that microinjection of oxytocin (OT) into the central nucleus of amygdala (CeA) induces hypergrooming in Wistar rats, a model of compulsion. The Wistar Audiogenic Rat (WAR) strain is a genetic model of generalized tonic-clonic seizures. Here we quantified grooming behavior in WAR, with *grooming scores*, flowcharts and directed graphs of syntactic and non-syntactic grooming chains, after bilateral administration of OT or saline (SAL) into the CeA and investigated the association between hypergrooming and imunohistochemistry of Fos activated compulsion networks and proposing a computational model of grooming behavior. The activated networks, driven from a CeA OT-dependent grooming pattern, in both Wistar and WAR were detected as Fos+ regions: orbitofrontal cortex, striatum, paraventricular nucleus of the hypothalamus, dentate gyrus, substantia nigra compacta and reticulata. In conclusion we can drive hypergrooming in WARs, defined previously as a model of ritualistic motor behavior in Wistar rats, with OT from CeA, a limbic structure and one of the principal amygdala complex outputs. Furthermore, our current pioneer behavioral and cellular description considers that hypergrooming (compulsion) in WARs is a comorbidity because: (1) WARs have the highest *grooming scores*, when exposed only to novelty (2) WARs have better *grooming scores* than Wistars after CeA-SAL, (3) WARs perform much better than Wistars in OT-CeA-dependent highly stereotyped behavioral sequences, detected by flowcharts as a combination of syntactic/non-syntactic grooming chains, (4) the behavioral sequences here demonstrated for grooming and hypergrooming can be modeled as quite reliable Markov chains and (5) with the exception of CeA-SAL injected animals, an exquisite map of brain Fos expression was detected in typical cortico-striatal-thalamic-basal ganglia-cortical circuit, among new areas, driven by OT-CeA.

**Author Summary:** Grooming is a complex set of regular behavioral sequences in rodents that can be mimicked with several pharmacological or molecular biology interventions. We have demonstrated previously that microinjection of the brain peptide oxytocin into the amygdala, a limbic region, induces hypergrooming in Wistar rats, a model of compulsion. The Wistar Audiogenic Rat strain is a genetic model of generalized seizures, in fact a model of epilepsy. Here we quantified grooming behavior in Wistar Audiogenic Rats, using several behavioral tools such as *grooming scores*, behavioral sequences and graphs of grooming chains, after bilateral administration of the oxytocin or its control (saline) into the amygdala. We also investigated the association between hypergrooming and activation of compulsion networks, proposing a computational (virtual) model of grooming behavior. Basically we were able to detect activated networks, driven from amygdala and the consequent oxytocin-dependent grooming pattern in epileptic and control animals. Those circuits are composed of cortical and subcortical areas, usually associated to the expression of motor rituals or compulsions. In conclusion, we can drive hypergrooming in epileptic animals, as compared to the previously defined model of ritualistic/compulsive motor behavior in control, rats. We concluded that hypergrooming (compulsion) is endogenously present in epileptic animals as a co-existent event (comorbidity), because when they were exposed to novelty, they express better *grooming scores* than control animals. The behavioral sequences here demonstrated for grooming and hypergrooming can be simulated as chains, where associations can be predicted from probabilities. Finally, an exquisite map of brain-activated cells was detected in both epileptic animal and their controls, in typical cortico-subcortical structures associated with rituals, but driven from a region which control emotions.

## 1. Introduction

Oxytocin (OT), a nonapeptide, synthetized in magno-and parvocellular neurons of the hypothalamic paraventricular (PVN) and supraoptic (SON) nuclei, is stored and released into the systemic circulation from the posterior pituitary [1]. PVN projects also to central targets [2] where OT receptors have been detected [1]. Central OT-ergic system is involved in grooming [3], maternal [4], sexual [5] and aggressive [6] behaviors, and has amnestic [7] antidepressant [8] and anxiolytic [5, 9] actions.

Obsessive-compulsive disorder (OCD) is characterized by obsessions (concerns, superstitious beliefs), and by compulsions (meaningless acts executed repeatedly) [10]. [11] detected a potential relationship between OT and OCD because patients with forms of OCD, without tics, had higher levels of cerebrospinal fluid OT than those with tics. Because of some discrepancies with [11], [12] suggested to study OT, in this context, as a brain peptide.

The Wistar Audiogenic Rat (WAR), strain, genetically selected for susceptibility to seizures [13,14], displays increased anxiety [15], and hyperactive hypothalamus-pituitary-adrenal (HPA) axis [16]. Therefore, because the WAR is an animal model of seizures with endogenous anxiety [15], and OT has anxiolytic effects [5, 9], we speculate that WAR OT-ergic system could be associated to anxiety and compulsion.

In that context, several behavioral alterations, are comorbidities occurring in patients with epilepsy, ranging from depression and anxiety to psychotic states [17,18]. In fact, there is a strong bidirectional connection between epilepsy and depression [19] and positive correlations between anxiety and epilepsy [17]. Moreover, amygdaloid complex functions include interpretation of emotional significance of sensory stimuli and appropriate responses [20]. Furthermore, the central nucleus of amygdala (CeA) participate in fear and anxiety responses Prefrontal cortex (PFC), striatum (ST), globus pallidus (GP) and thalamus (TH) constitute a neural network involved in switching behavior patterns [21]. Anatomical and functionally distinct areas of the cortex, TH and basal ganglia (BG) are connected by segregated and parallel circuits that originate in a particular cortical area, while ST regions receive projections from cortical areas and send them to GP, SNPr and TH; a dysfunction in this network could explain the pathophysiology underlying OCD [22].

We specifically aimed to characterize the activation of a circuitry related to compulsive grooming induced by OT in naïve (no seizures experience) WARs, an experimental model of seizures [13, 14], as compared to the one we already demonstrated in Wistars [23], using behavioral tools such as scores [24], flowcharts [25] and graphs [26] of syntactic and non-syntactic chains [27, 28], and cellular tools such as OT and c-Fos microscopy with the aim of building a Markov chain model [29, 30]. The interpretation of the data was done in the context of the pathophysiology of the comorbidity of epilepsy and compulsion.

## 2. Materials and Methods

### 2.1. Animals

Wistar rats (n=11) and WARs (n=18), with 250 - 280 g at the beginning of the experiments, from the Animal Facilities at the Campus of Ribeirão Preto, University of São Paulo, Brazil, were housed under controlled conditions and maintained on a 12 h light/dark cycle with ad-libitum access to food and water.

Animals were divided in groups microinjected bilaterally into CeA, being WAR-OT (n=6), WAR-SAL (n=6); Wistar-OT (n=6), Wistar-SAL (n=5). An additional non-injected (naïve) group WAR/Basal (n=6) was used for behavioral analysis only. For statistical purposes, only animals that were correctly implanted bilaterally into CeA were used. All procedures were approved by the Ethics Committee on Animal Experimentation of the Ribeirão Preto School of Medicine - University of São Paulo (protocols 054/2007 and 151/2011).

### 2.2. Surgery

Rats were anesthetized with ketamine (1,0 mg/kg i.p.; União Quiímica Farmacêutica Nacional, Embu-Guaçu, Brazil) and xylazin (0,7 mg/kg i.p.; Bayer, São Paulo, Brazil) and bilaterally implanted into CeA, according to [53] coordinates: antero-posterior: −2.0 mm (bregma); medio-lateral: 4.0 mm and dorso-ventral: 7.0 mm.

### 2.3. Microinjection procedures into CeA

Animals were handled during three days and seven days after surgery, received either injection of sterile SAL (NaCl 0,9 %) or OT (0,5 μg in saline; Sigma). Drugs were delivered into CeA with a 5 μl syringe (Hamilton, model 95N) connected to a polyethylene tubing (PE-10) with an infusion pump (Harvard Apparatus - PHD 2000), 200 nl of SAL or 200 nl of OT. The needle was maintained 1 minute inside the guide cannula after the infusion, when the rats were placed in an acrylic box (30 × 30 × 30 cm) and their behavior was video recorded. All injections were done between 16:00 h and 18:00 h.

### 2.4. Behavioral analysis

Recordings (1 hour) began with the animal placed in an acrylic box after OT or SAL CeA microinjections. See details for the calculation of the grooming score [24] in Fig. 1.

**Figure 1.**
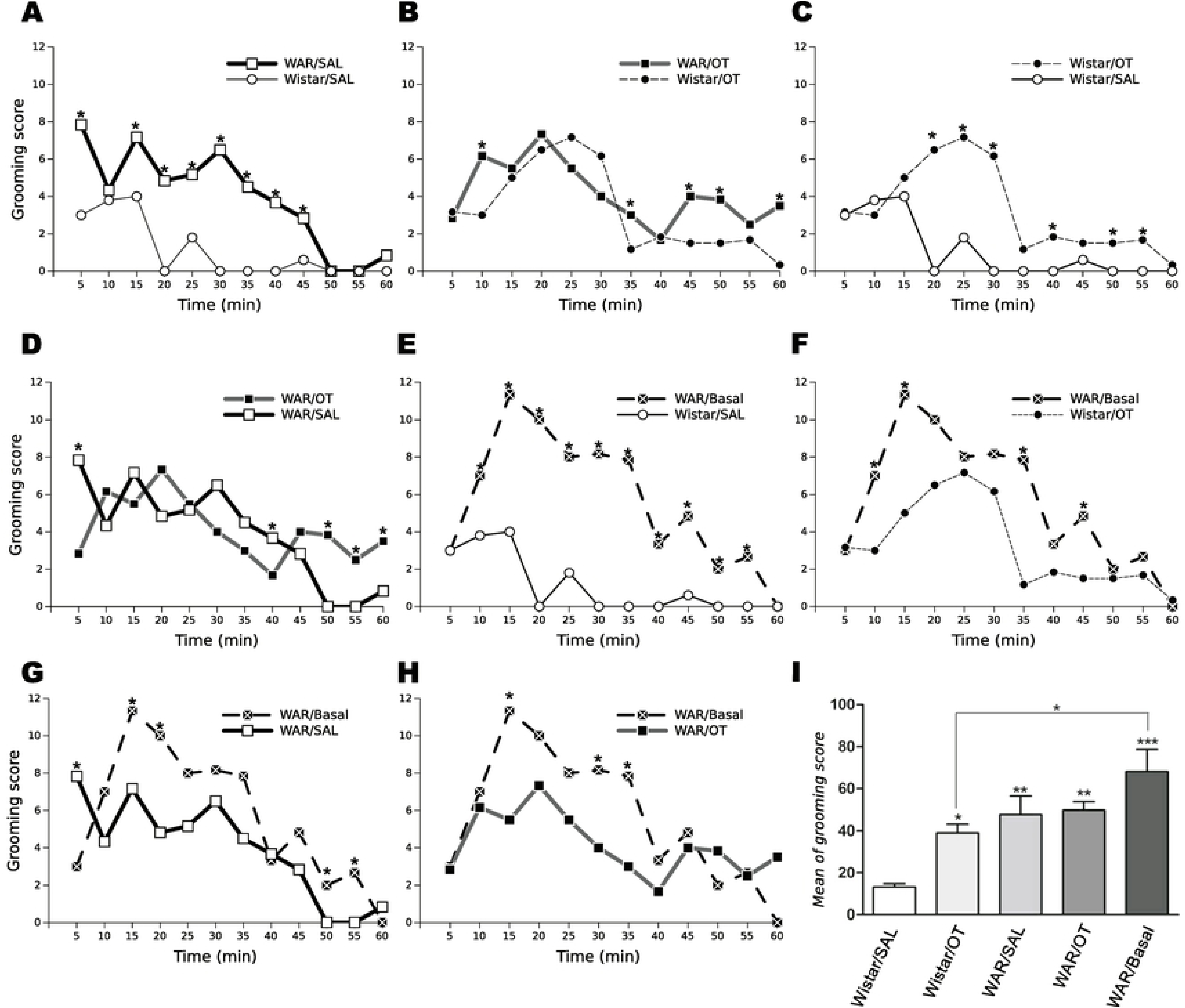
Time course of grooming behavior. Every 15 seconds, a point was scored if the animal displayed one of the following events: washing, general grooming, scratching, licking of paws, grooming of head, grooming of face, grooming of anogenital region or tail, or grooming of another region of the body. The sum of points in 60 minutes is called the grooming score, as described by (Gispen et al., 1975), and the maximum value was 240 points. Illustrated are the grooming scores of animals of the groups Wistar/SAL, Wistar/OT, WAR/SAL, WAR/OT and the group WAR/Basal after bilateral microinjections into CeA. Time course of the effect of SAL or OT injection on grooming behavior. (*p < 0.05; one-way ANOVA with repeated measures followed by pairwise comparison with Bonferroni correction). When compared the two strains with the same drug (WAR/SAL vs. Wistar/SAL) it was observed higher frequency of grooming during the 60 minutes in the group WAR/SAL (A). After injection of OT, the frequency of grooming was higher in the WAR/OT compared with Wistar/OT in the whole analyzed time window (B). In the groups of same strain with different drugs, the frequency of grooming was higher in the Wistar/OT when compared with Wistar/ SAL in the observation windows of 15-20, 20-25, 25-30, 35-40, 45-50 and 50-55 minutes (C). The frequency of grooming behavior was higher in the WAR/OT when compared with the group WAR/SAL at intervals of, 45-50, 50-55 and 55-60 minutes. Opposite occurred between 0-05, 35-40 minutes (D). All groups Wistar/SAL, WAR/SAL, Wistar/OT and WAR/OT compared with WAR/Basal: grooming scores reached the greatest expression in the WAR/basal (E, F, G, H). Comparing both strains with OT, the total mean of grooming score in the group WAR/OT was higher than Wistar/OT (I). Grooming score average count of groups of animals injected bilaterally with SAL or OT in CeA: Wistar/SAL, Wistar/OT, WAR/SAL, WAR/OT and the group WAR/Basal. (*p < 0.05, **p < 0.001, ***p < 0.0001; one-way ANOVA followed by pairwise comparison with Bonferroni correction).

Besides the grooming score, we used flowcharts, statistical associations between pairs of behaviors (dyads) (see calibration in Fig. 2A) [23, 25], and directed graph analysis, visualization of syntactic and non-syntactic grooming chains [27, 28], which measure behavior complexity [26].

**Figure 2.**
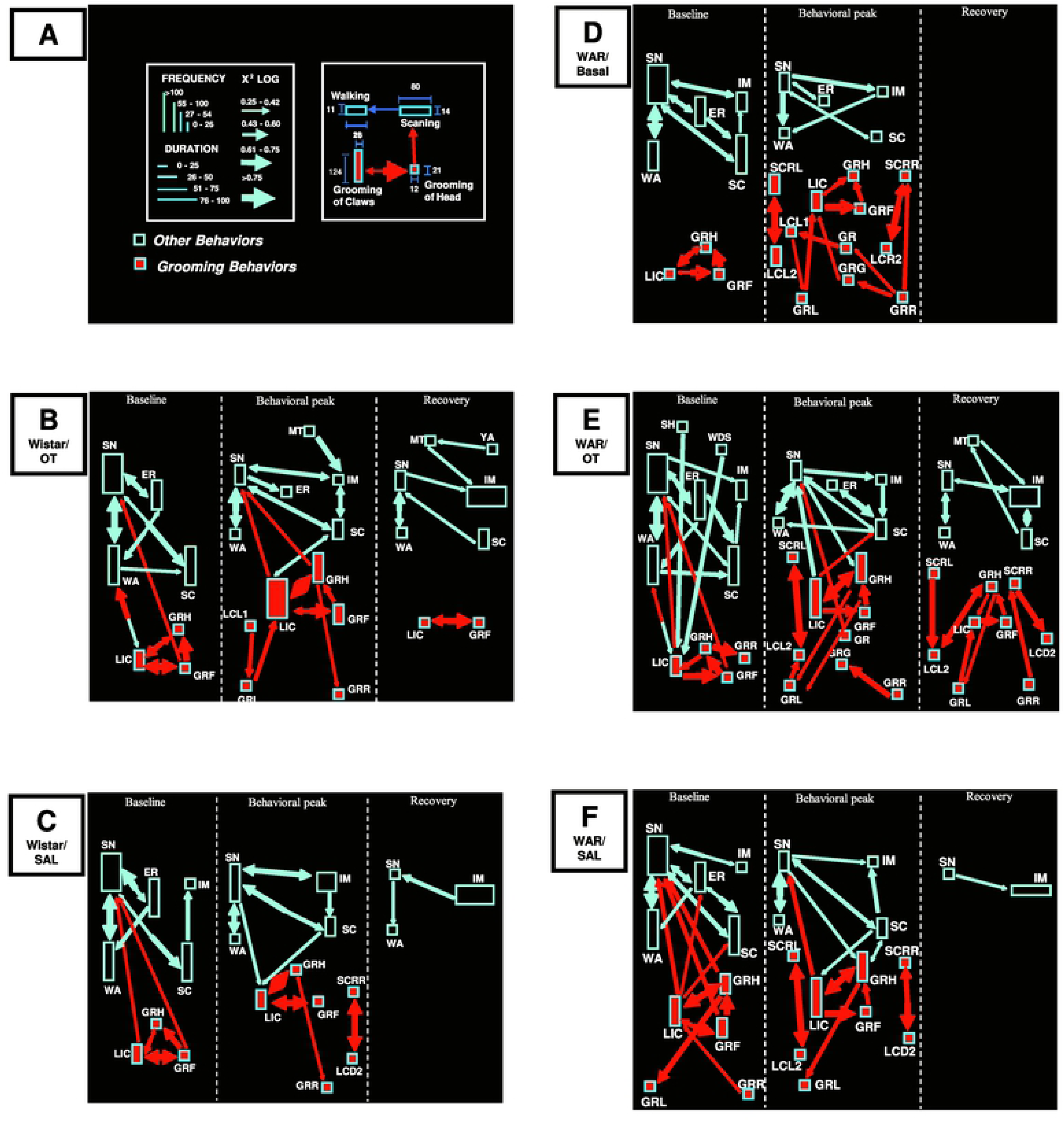
Neuroethological evaluation of behavioral sequences after SAL or OT bilateral microinjections into the CeA. Baseline Period, Behavioral Peak Period and Recovery Period are flowcharts from 3 minutes observation windows. (A) Flowchart calibration: the height of rectangles represents the frequency of a behavioral item and the base corresponds to the duration during the observation windows; the arrows represent statistical values (χ^2^ > 3.84; P < 0.05) highlighting the strength of association between pairs of behaviors (dyads). The colors identify the main behavioral clusters. The use of colors for clusters in the flowcharts is for illustration purposes and does not have any influence in the statistical analysis. An interaction between the grooming behavioral cluster and the exploratory activity cluster is observed in groups Wistar/OT (B), Wistar/SAL (C), WAR/OT (E) and WAR/SAL (F) which does not occur with the group WAR/Basal (D). We also observed that in the Behavioral Peak Period there was mainly a decrease in the frequency of exploratory behaviors compared with the Baseline Period. An interesting category of behaviors involving SCRL and LCL2 (left side grooming) appeared only, with strong significance, in the WAR strain (D, E and F). In the Recovery Period, only Wistar-OT and WAR-OT groups showed grooming. In the group Wistar/OT, we observed an interaction only between LIC and GRF (B), which diverges from WAR/OT (E), where several items of the grooming cluster interacted among themselves. In the Recovery Period, the group WAR/Basal (D) displayed no behavioral interactions between the exploratory activity cluster and the grooming cluster; the WAR/SAL (F) showed only interaction between SNF and IM while WAR/OT (E) has significant interactions involving five exploratory behaviors. The group Wistar/SAL has only interactions between exploratory behaviors WA, SNF and IM, whereas Wistar/OT exploration activity involved interactions with six exploratory behaviors. (B) Group Wistar/OT, (C) Group Wistar/SAL, (D) Group WAR/Basal, (E) Group WAR/OT and (F) WAR/SAL. **Grooming Behaviors**: **GRR** - Grooming of body (right); **GRL** - Grooming of body (left); **GRG** - Grooming of genitalia; **GRH** - Grooming of Head; **LIC** - Licking of Claws; **LCR1** - Licking of Claws (right, anterior); **LCR2** - Licking of Claws (right, posterior); **LCL1** - Licking of Claws (left, anterior); **LCL2** - Licking of Claws (left, posterior); **GRF** - Grooming of Face; **SCRL** - scratch left, **SCRR** - scratch right. Exploratory and other behaviors: **ER**-Erect Posture; **IM** - Immobility; **MT** - Mastigatory Movements; **PIV1** - Pivoting (movement of head, laterally); **SCRR** - Scratching of body (right); **SCRL** - Scratching of body (left); **SH** - Head Shaking; **SCA** - Scanning; **SNF** - Sniffing; **WA** - Walking; **WDS** - Wet Dog Shaking; **YA** – Yawn; Grooming Behaviors: **GRR** - Grooming of body (right); **GRL** - Grooming of body (left); **GRG** - Grooming of genitalia; **GRH** - Grooming of Head; **LIC** - Licking of Claws; **LCR1** - Licking of Claws (right, anterior); **LCR2** - Licking of Claws (right, posterior); **LCL1** - Licking of Claws (left, anterior); **LCL2** - Licking of Claws (left, posterior); **GRF** - Grooming of Face; **SCRL** - scratch left, **SCRR** - scratch right.

Recorded sessions were analyzed with a behavioral dictionary (codes in Figure 2 legend). Three periods were evaluated: the Baseline Period, represents the initial 3 minutes (0-3 minutes); the Behavioral Peak Period, exacerbation of grooming behavior, varying among the groups; and the Recovery Period, final 3 minutes of observation (57-60 minutes).

Directed graphs were built from the Behavioral Peak Period, with each behavioral category considered a node, connected by an edge with the following behavior. The position of each node depends on the phase to which this behavior belongs, following the IV phases of syntactic grooming chains: I, elliptical strokes, II, unilateral strokes, III, bilateral strokes, and IV, body licking [27, 28]. Therefore, we analyzed whether during the grooming Behavioral Peak Period the rats develop the IV phases of syntactic chains in sequential order and the time spent in each one, or a different grooming pattern for each of the experimental groups, estimating, as indicated in the legend of Figure 3 the complexity measures of the node of the directed graph [26], also, using the Erdős–Rényi model [32], checking if the directed graphs were random or not.

**Figure 3.**
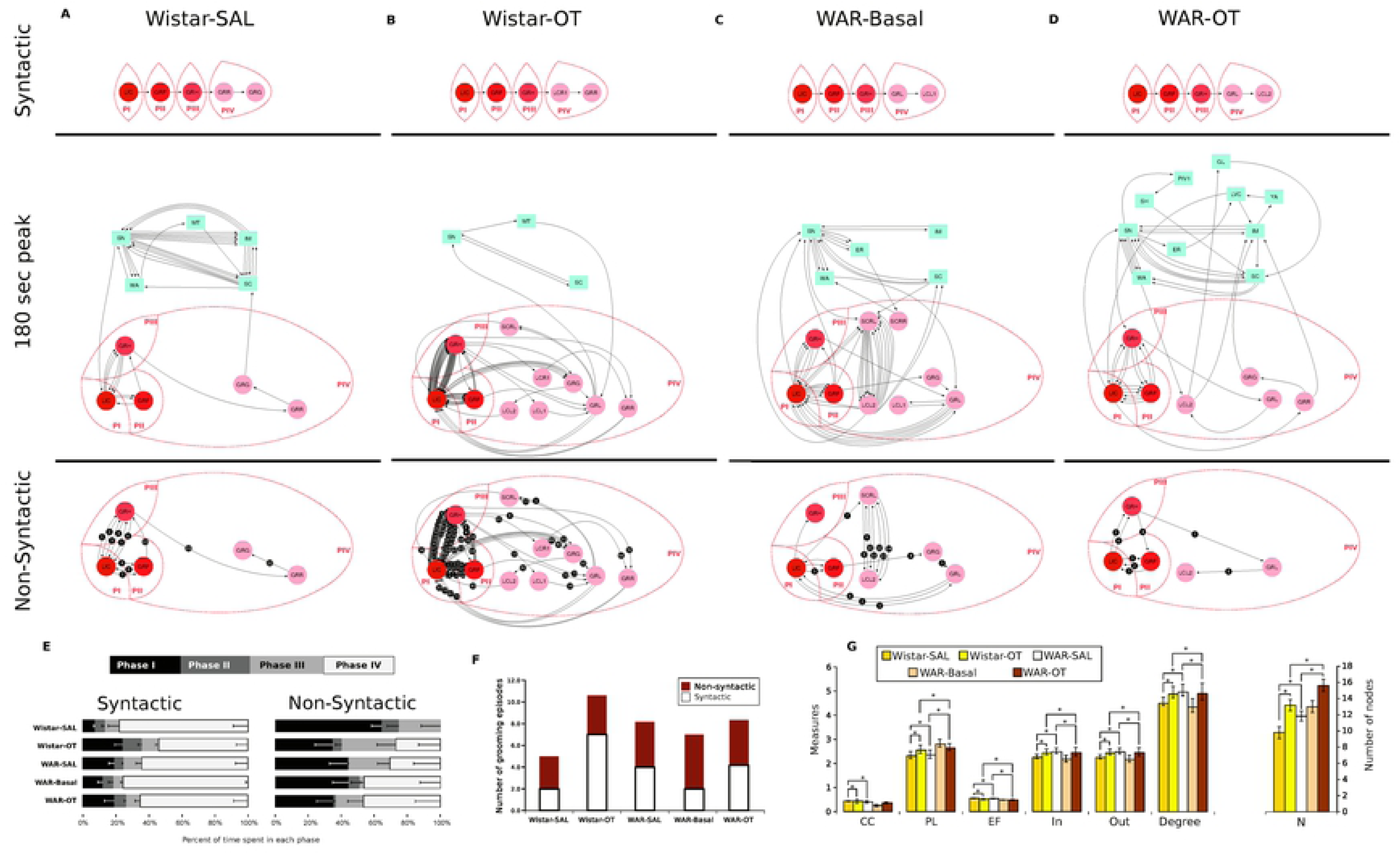
Directed graph of the peak of grooming from which were extracted both syntactic and non-syntactic grooming chains from individual examples of each of the following groups: A) Wistar/SAL, B) Wistar/OT, C) WAR/Basal D) WAR/OT. E) Average time spent in each one of the phases depends on the existence of either syntactic or non-syntactic grooming chains for each one of the experimental groups. Error bars represent standard error. F) Number of sequences of the syntactic and non-syntactic grooming chains during the peak of grooming in the five groups. G) Average Cluster coefficient (CC), Path length (PL), Efficiency (EF), in-degree (In), out-degree (Out) and total: (in-degree+out-degree) (Degree), estimated from directed graphs of the peak of grooming in the five groups. Error bars represent standard error and asterisk represents significant difference (* p < 0.05; independent t-test with Bonferroni correction for multiple comparisons). Grooming Behavior: **Phase I**: **LIC**-Licking of Claws; **Phase II**: **GRF**-Grooming of Face; **Phase III**: **GRH**-Grooming of Head; **Phase IV**: **GRB**-Grooming of body, **GRL**-Grooming of body (left), **GRR**-Grooming of body (right), **GRG**-Grooming of genitalia, **LCL1**-Licking of Claws (left, anterior), **LCL2**-Licking of Claws (left, posterior), **LCR1**-Licking of Claws (right, anterior), **SCRL**-scratch left, **SCRR**-scratch right.

Based on the directed graph, Markov chain models were built, in which each node represents the chain states. The probability of transitions, from one state to another, were estimated considering all behaviors of a specific group in Syntactic and Non-Syntactic grooming chains. The Markov chains properties were evaluated with the aim of validating the hypothesis of grooming chains being stochastic processes of reduced memory. This particular condition expands the possibilities of mathematical characterization of this kind of behavior.

### 2.5. Perfusion, tissue preparation and immunohistochemistry

Animals were perfused 90 minutes after OT-CeA microinjection to evaluate c-Fos expression [33]. Briefly, animals were deeply anesthetized with sodium thiopental (Abbott, North Chicago, USA) 60 mg/Kg and transcardially perfused with 100 ml of saline phosphate buffer 0.01M (PBS pH 7.4), followed by 300 ml of 4% paraformaldehyde (PFA; Acros Organics, New Jersey, USA).

Brains were post-fixed for about two hours in PFA 4%, cryoprotected in 20% sucrose for 1-2 days at 4 o C before being frozen in dry ice-cold isopentane and sectioned in 40 m-thick coronal sections on a cryostat (Micron-Zeiss HM-505-E; Walldorf, Germany). Sections were mounted on gelatin-coated slides and stained with cresyl violet to check cannula location.

For immunohistochemistry (OT) and immunofluorescence (OT, c-Fos), assays followed [23] and [34]. Briefly, antibodies dilution and incubation times were: goat anti c-Fos (1:500; Santa Cruz Biotechnology INC.; CA, USA; 1:500), 20 h in wet hood at room temperature. Rabbit anti OT (1:2000; Peninsula Laboratories, INC, Belmont, CA, USA), 20 h in wet hood at room temperature.; Alexa 488 anti-goat (1:1000; Molecular Probes; Oregon, USA), 1 h at room temperature in wet hood and IgG biotinylated rabbit anti goat (1:500; Calbiochem) two h at room temperature in wet hood). For immunohistochemistry staining we used DAB-peroxidase reaction with Vector Vectastain Elite ABC kit (pk-6100) and Vector peroxidase substrate kit (sk-4100).

### 2.6. Acquisition and analysis of histology images

All the sections/images were captured/analyzed with a digital system composed of an Olympus BX-60 (Tokyo, Japan) microscope, a CCD-Cool camera (Optronics 750 L, Goleta, CA, USA) and the Image-Pro-Plus software (Media Cybernetics).

### 2.7. Cell counting

Initially, using a semi-quantitative scale, histological sections containing each entire nucleus were prepared and 6 sections per nucleus were selected for bilateral counting. Immunohistochemically c-Fos+ cells were searched into the pre-frontal cortex (PFC) (AP: 4.20 mm to 3.20 mm), ST (AP: −1.20 mm to −1.90 mm), DG (AP: −1.80 mm to −2.80 mm), substantia nigra compacta (SNPc) and SNPr (AP: −5.20 mm to −6.30 mm), according to the stereotaxic coordinates of [31]. In a second counting analysis several of the sections containing the PVN were prepared and three sections were selected for counting: anterior (AP:-1.40 mm) middle (AP:-1.80 mm) and posterior (AP:-2.12 mm). The neurons labeled c-Fos+ and OT+ (co-localization) in the PVN were counted using ImageJ (Wayne Rasband; Research Services Branch, Bethesda, Maryland, USA)

### 2.8. Statistical analysis

For grooming scores we used a Poisson model and PROC NLMIXED (SAS) [23, 56]. For flowcharts see [25, 36] and Results Section. For the graphs, we used analysis of variance one-way NOVA, followed by pairwise comparison with Bonferroni correction and χ2 independence test to compare the syntax of the grooming chains. Finally, for the neuroanatomical data and the average of grooming score, we used one-way ANOVA followed by post-hoc Newman-Keuls test. The significance used in all tests was P < 0.05 and we used GraphPad Prism or R statistical programming language.

## 3. Results

### 3.1. Behavior

The time course of grooming behavior added in intervals of 5 minutes (60 total) is shown in Fig. 1(A-I), comparing WAR/SAL vs. Wistar/SAL, (B) WAR/OT vs. Wistar/OT, (C) Wistar/SAL vs. Wistar/OT, (D) WAR/OT vs. WAR/SAL, (E) Wistar/SAL vs. WAR/Basal, (F) Wistar/OT vs. WAR/Basal, (G) WAR/SAL vs. WAR/Basal and (H) WAR/OT vs. WAR/Basal.

### 3.2. Flowcharts

All behavioral sequences are illustrated in Fig. 2(B-F). At the *Baseline Period* of each group, we observed a cluster of behaviors involving GRH, LIC and GRF with interactions among all items. The WAR/OT group has also interactions with GRR which follows GRH (Fig. 2, E). Wistar/SAL presents GRL, GRR and GRH that follow LIC (Fig. 2, C), similar to all groups, except for WAR/Basal (Fig. 2, D). All other groups display significant interactions between the grooming behavior cluster and the exploratory activity cluster. At the *Behavioral Peak Period*, we observed a higher frequency of behaviors such as GRH, LIC and GRF, when compared with the *Baseline Period*. Moreover, the WAR strain has a greater expression of behaviors (Fig. 2, D, E and F), strongly supportive of OT-CeA-induced and endogenous (WAR-Basal) *hypergrooming*. See additional description of behavioral sequences in Fig. 2.

### 3.3. Syntactic and Non-Syntactic Grooming Chains

All experimental groups showed both syntactic and non-syntactic chains, and the most complex behavioral sequences, when compared to Wistar/SAL (Fig. 3A) were observed in Wistar/OT, WAR/OT and WAR/Basal (Fig. 3B, C and D, respectively).

During the grooming sequences with syntactic chains, the rats of all of groups spent more time in P IV than in the other phases (F_(3,72)_ = 20.173, p -value < 0.05)(Fig. 3E). This was different from the grooming sequences with non-syntactic chains, where we found differences (F_(3,112)_ = 6.0073, p-value < 0.05) between time spent on P I, P II and P III, and also between P II and P IV for all of the groups. (Fig. 3E). The number of sequences of the syntactic and non-syntactic grooming chains during the *Behavioral Peak Period* did not show significant differences between the groups (χ^2^_(4)_= 2.5659, p > 0.05)(Fig. 3F). Regarding the directed graphs, we only found differences in the PL of grooming sequences with and without syntactic chains. The sequences with syntactic chains presented bigger PL (F_(9,12.03)_ = 14.935, p < 0.05) for almost all the groups, with exception of Wistar/SAL. On the other hand, from the ErdösRényi coefficient [32], the directed graphs of the grooming chain are not random, with a maximum likelihood estimator of the ErdösRényi model [32] of −0.6748.

Additionally, the directed graphs of the *Behavioral Peak Period* showed differences between Wistar/SAL and all the WAR group for all of the measures (CC, PL, EF, In, Out and Degree [total]). Also, qualitatively, the directed graph of the *Behavioral Peak Period* of the WAR groups showed higher number of exploratory behaviors and, as well, more interaction between grooming and exploratory behaviors than the Wistar groups.

### 3.4. Localization of Fos Immunohistochemistry

The Fos expression map associated to OT-CeA induced grooming is illustrated in Fig. 4 for the groups: Wistar/SAL, Wistar/OT, WAR/SAL and WAR/OT.

**Figure 4.**
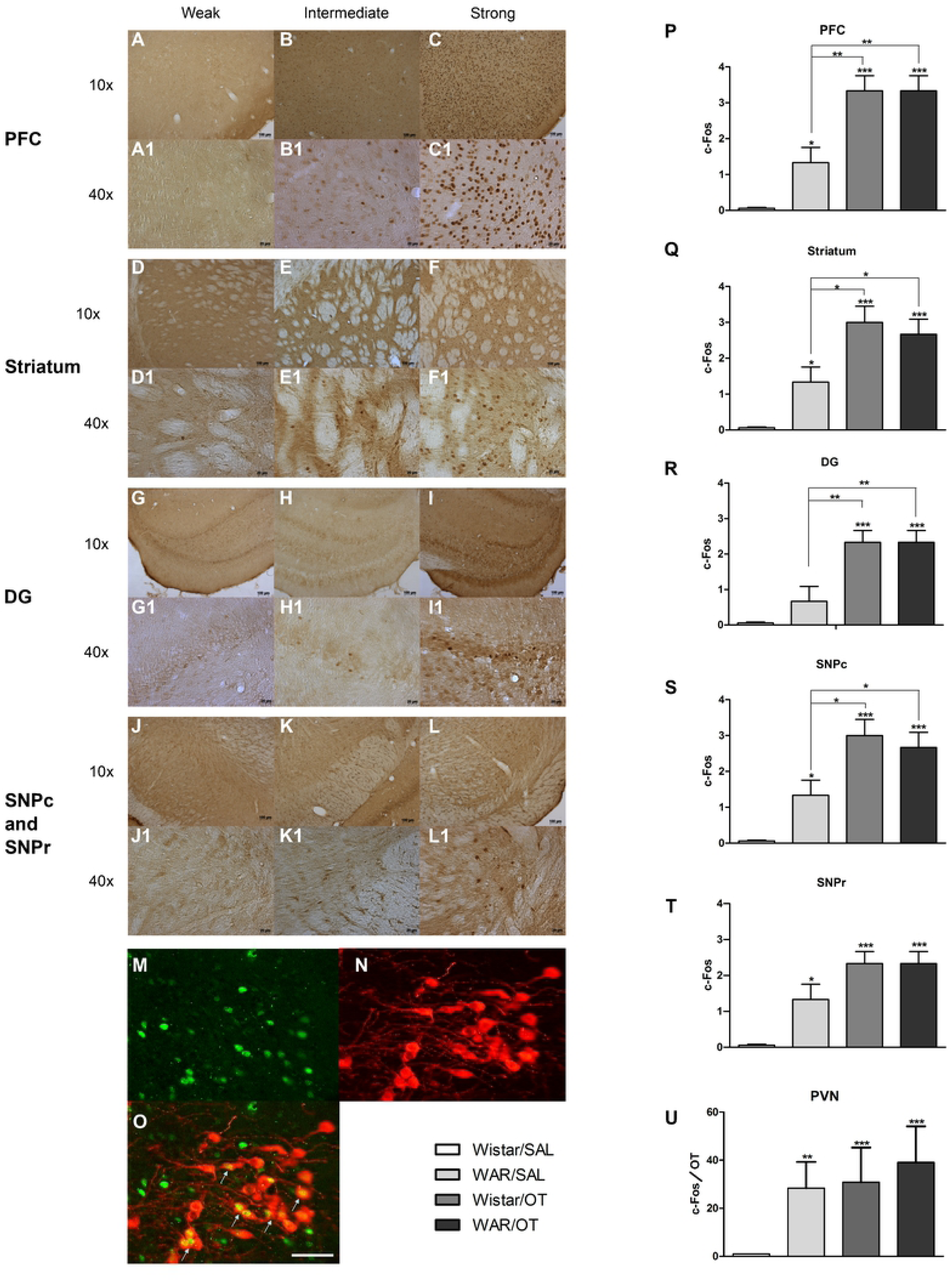
Mapping OT and Fos distribution. In the Fos quantification scale, 0= low labeling; 2=intermediate labeling and 4= high level labeling, using as reference the intensity of labeling and number of neurons. Photomicrographs showing Fos+ neurons in PFC **(A)** and (A1-zoom) indicating low labeling **(B)** and (B1-zoom) intermediate labeling and **(C)** and (C1-zoom) high labeling, Striatum in **(D)** and (D1-zoom) low labeling **(E)** and (E1-zoom) intermediate labeling and **(F)** and (F1-zoom) high labeling, DG in **(G)** and (G1-zoom) indicating low labeling **(H)** and (H1-zoom) intermediate-labeling and **(I)** and (I1-zoom) high labeling, SNPc and SNPr in **(J)** and (J1-zoom) low labeling **(K)** and (K1-zoom) intermediate labeling and **(L)** and (L1-zoom) high labeling. Photomicrographs showing co-localization of OT (red) and Fos (green) in the PVN. **(M)** OT, **(N)** Fos and **(O)** co-localization of OT/ c-Fos (represented by arrows). Fos+ cell counts in the PFC **(P)**, Striatum **(Q)**, DG **(R)**, SNPc **(S)** and SNPr **(T)** in the 4 experimental groups. * p < 0.05, ** p < 0.001, *** p < 0.0001. **(U)** Counts of co-localized neurons containing OT+ and Fos+ in the PVN of the four experimental groups. ** p < 0.001, *** p < 0.0001. Briefly, Fos+ cells were detected in PFC **(A-C1)**, ST B, DG **(G-I1)** SNPc and SNPr **(J-L1)**. Counting of Fos+ neurons had significant differences in the groups Wistar/OT, WAR/SAL and WAR/OT, when compared with Wistar/SAL **(P-S)**. Different strains with the same drug, for example, Wistar/OT and WAR/OT had no differences in the expression of Fos+ cells. Groups with OT compared with WAR/SAL presented greater expression of Fos+ neurons particularly in PFC, DG, ST, SNPc after OT-CeA. Scale bar 250 μm.

### 3.5. Co-localization of OT and Fos immunofluorescence

We found wide co-localization of OT+/Fos+ immunofluorescent neurons in the PVN (Fig. 4M-O), consistent with the expression of the grooming data after OT-CeA. In fact, only the group Wistar/SAL present lower behavioral activity (Fig. 1E and I) and lower Fos+ activation of PVN in the neuroanatomical (Fig. 4U) analysis.

### 3.6. Behavioral expression of [OT-CeA]-driven grooming behavior and proposed neural networks

In Figure 5 are illustrated comparative graphs of the *Behavioral Peak Period* of grooming of all animals from the Wistar-OT and WAR-Basal groups and the complementary addition of traditional neural networks involved with OCD and those proposed after current data, consequent to Fos+ activation of several nuclei and their connections after OT-CeA.

**Figure 5.**
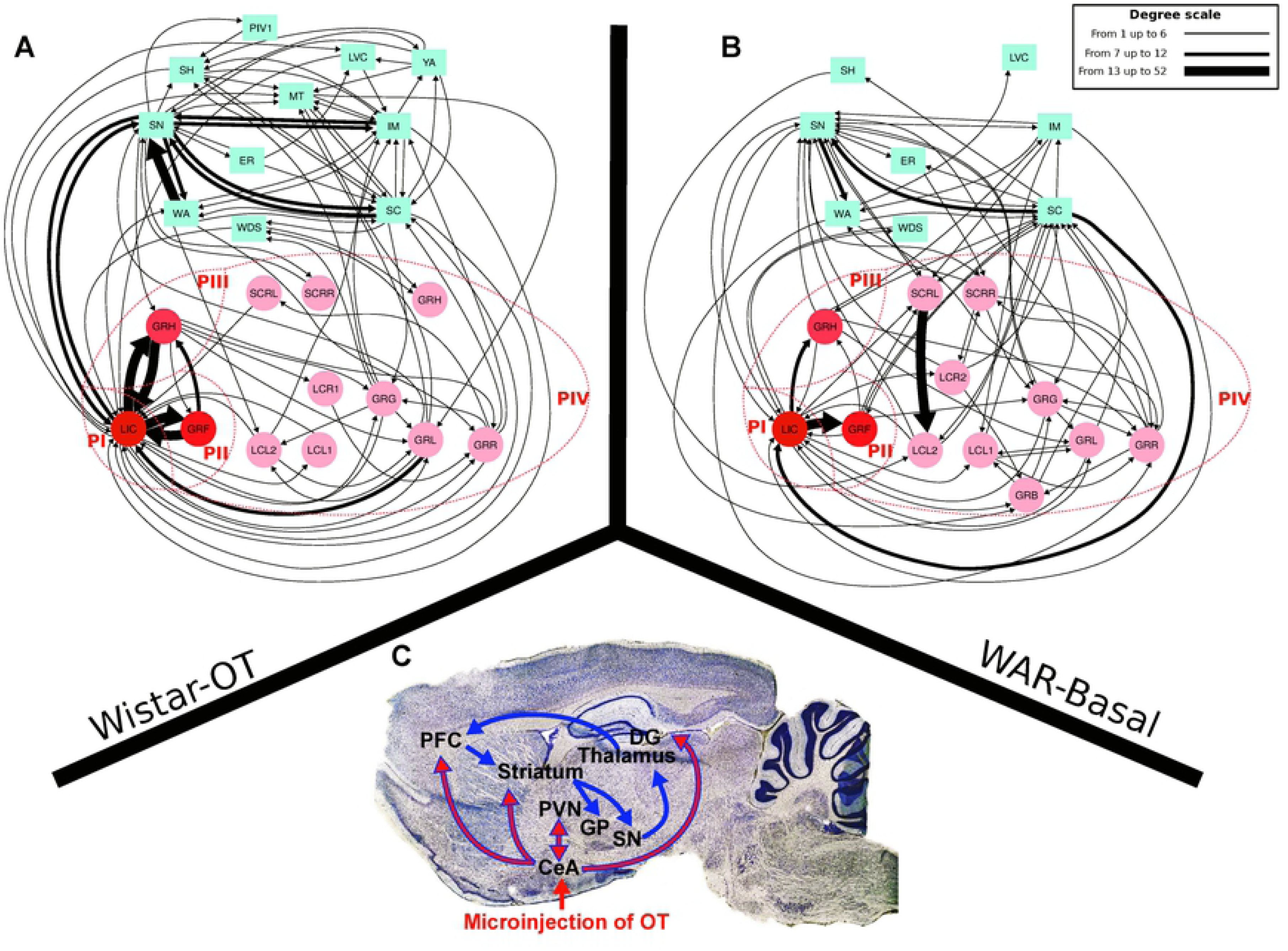
Directed graph of the peak of grooming of all animals from the Wistar-OT **(A)** and WAR-Basal **(B)** groups. The edges represent different numbers of connections (see degree scale). **(C)**, Photomicrograph of a sagittal section of the brain of one rat, removed from the atlas of Paxinos and Watson (2005) showing a neural network involved in switching behav-ioral patterns of OCD described in the literature in blue arrows and in red arrows showing Fos+ (activated) nuclei and their connections after the current microinjections of OT in CeA. Briefly, current behavioral and Fos data amplified the neural networks associated to the expression of hypergrooming as a compulsive behavior, driven by OT-CeA. In addition to the traditional cortico-striatal-thalamic-cortical network (blue arrows), limbic (CeA) driven activity (red arrows) recruits, PVN, DG and PFC. Both kinds of hypergrooming, the OT-CeA driven and the one induced by novelty in WARs, are proofs of this compulsion as a comorbidity in a strain which is already genetically susceptible to epilepsy. For additional functional neuroanatomical background see Graybiel and Saka (2002) on basal ganglia and Kalueff et al. (2016) for complexity of neurobiology of grooming behavior.

### 3.7. Markov chain model

The syntactic and non-syntactic grooming sequences fit well with Markov chain properties, evaluated using a Chi-square test. The transition probability matrices (see Fig. 6) reproduce properly the sequence of events performed by the animal of a specific group, such as WAR/OT Non-Syntactic chain.

**Figure 6.**
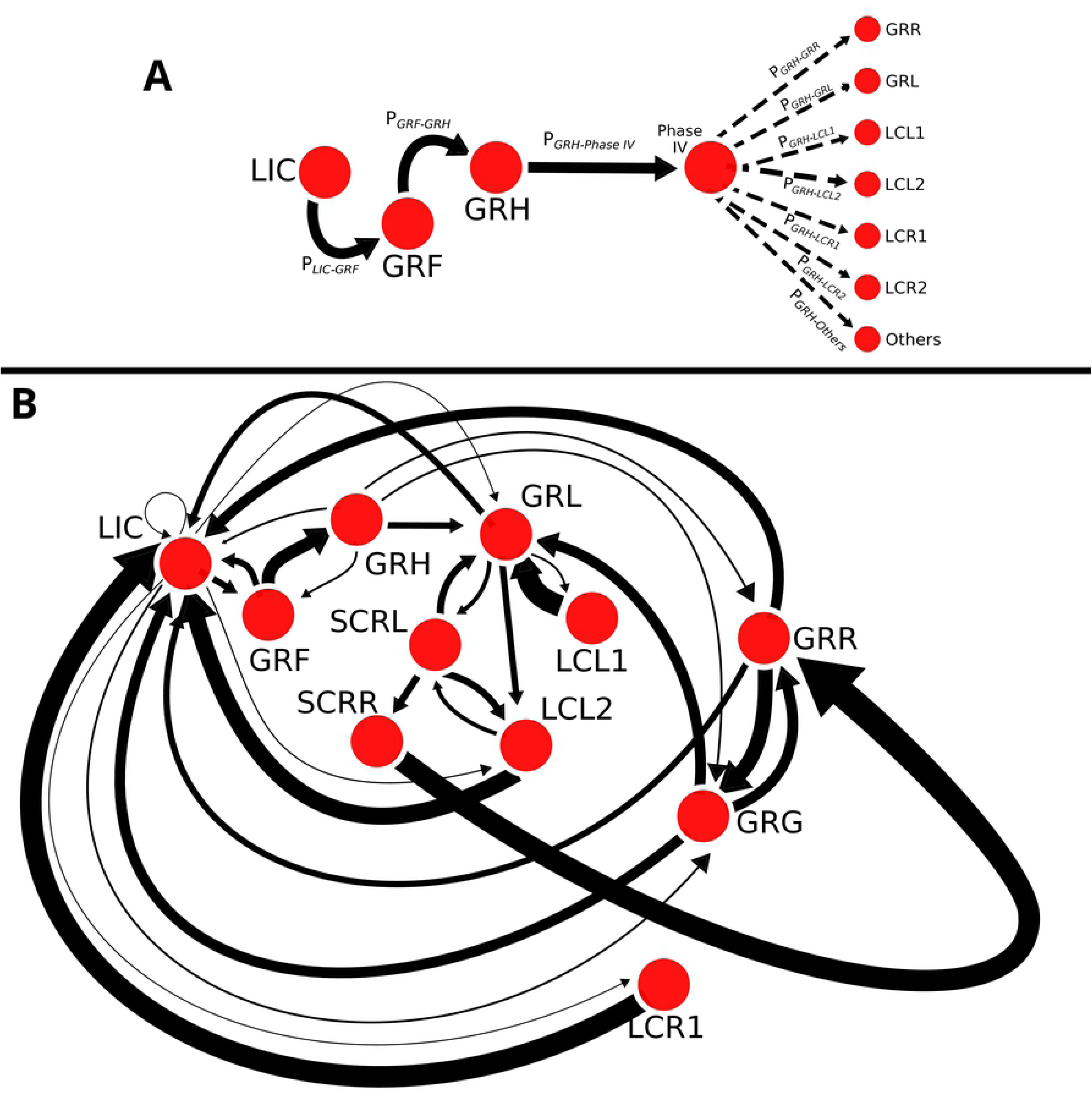
(**A**) Graph representation of the generic Markov chain model for a grooming episode, with the behaviors in the nodes and the probability to change from one behavior to another in the edges. (**B**) The model of the non-syntactic grooming episode for the WAR-OT group. The thickness of the edges represents the probability of transition from one state to another.

## 4. Discussion

The mean *grooming score* after microinjection of SAL or OT in the CeA of the WAR strain clearly showed hypergrooming, in agreement with our experimental model of compulsion in Wistars [23], and the fact that WARs are endogenously anxious [15] and have a hyperactive HPA system [16].

*Grooming score analysis* demonstrates that WARs maintain a sustained compulsive behavior, because, while Wistar-OT showed a period of maximum activity between 20-30 minutes, grooming behavior still stayed exacerbated in WARs 60 minutes after CeA-OT, suggesting an endogenously activated circuit. Interestingly OT-dependent grooming in Wistars [23], induces cardio-respiratory alterations, lasting near forty minutes [37]. Therefore, having a model in which we can monitor cardio-respiratory responses, during the motor expression of compulsions, is of enormous importance for patients with OCD, because it is well known that intrusive thoughts (obsessions), induce anxiety, and compulsive rituals have anxiolytic effects [38], the anxiety reduction theory of OCD [39].

OT-CeA or SAL-CeA in Wistar rats indicates that up to the first 5 minutes the grooming behavior was the same, revealing the stress evoked by exposure of the animals to novelty. In the case of the WAR-Basal, the increased grooming in the absence of drugs is a strong proof of endogenous compulsive grooming (comorbidity).

WAR/OT-CeA (contrasting to WAR/-SAL) shows an exacerbation of grooming behavior, up to 35 minutes, after which they maintained the behavior, suggesting the influence of OT-CeA in a modulatory circuit of motivation [40] associated to compulsions. Another important finding is that in the *Recovery Period*, WAR animals microinjected with OT displayed strong interactions between SCRL, LCL2, GRH, LIC, GRL, GRF SCRR, GRR and LCR2, reflecting hypergrooming.

Grooming behavior is generally initiated by paw licking or face-washing, followed by grooming of the fur around the head, neck, and body in a cephalocaudal stepwise pattern [27,41]; one particular type of cephalocaudal ordering is the “syntactic chain” sequence [27,28]. The current graph analysis [26] evaluates continuously behavior, its quality and complexity. The meaning of behavioral chains is not extracted from frequency or duration of their elements (as in flowcharts), but from their ordering. In that sense, OT-CeA grooming or the one in the WAR/Basal group, are not similar to grooming occurring after SAL. A variety of sequencing rules appear to be “*obeyed by”* the nervous system and expressed as grooming [28]. The information of directed graphs of the syntactic grooming chains demonstrates that these actions are not randomly expressed but instead, have marked sequential dependency, demonstrated with the negative value of the maximum likelihood estimator of the ErdösRényi [32].

The directed graph also showed that, for almost all of the groups, nonsyntactic grooming chains evolve more frequently and longer from PI to PIII. In contrast, at least qualitatively, non-syntactic chains of the WAR groups look simpler than Wistar/OT chains, because they were more occurrences of exploratory behaviors that interrupt WAR non-syntactic chains.

Additionally, the comparison of the directed graph of the *Behavioral Peak Period* allows identifying fine details of grooming behavior that clearly differentiate patterns in the Wistar/SAL, with exceptional transitions from grooming to exploratory behaviors, from the other groups. Those graphs allow also differentiating between Wistar/OT and WAR groups, including WAR-Basal, because in the latter there is more prominent occurrence and interactions between grooming and exploratory behaviors.

The Markov chain models of syntactic and non-syntactic grooming chains (see Fig 6) suggest that these episodes can be correctly described by memory-one models, which make possible the use of the range of measures available for these kinds of models, such as *mean recurrence time, expected number of visits, absorbing states, among others*, extending the mathematical characterization with measures which, not only describe, but can predict the grooming episode for a specific condition. For instance, the estimation of the absorbing states could identify the behaviors in which the grooming episode ends, offering insights regarding the neuronal activity in which the loop of circuit activation, which provoked the grooming episode, stopped. Other measures, such as mean recurrence time, might offer information about the proportion of time that the animals spend making a specific behavior, and in this way, improving the behavioral characterization [30,42].

Now, what are the brain Fos+ networks that are activated by OT-CeA, accompanying compulsive grooming behavior? How can CeA (OT injection site) and the PVN (OT origin site), in fact part of the hypothalamic grooming area (HGA), coordinate the expression of compulsive grooming behavior? We presently found Fos+ neurons, activated by the OT-CeA/grooming stimulus, in animals of both WAR and Wistar strains, in addition to classical OCD areas in PVN, PFC, ST, DG, SNPc and SNPr. As per the co-localization of OT and Fos+ neurons in the PVN, we verified that all groups showed a statistically significant difference, when compared to the Wistar/SAL group and that the WAR strain displays an endogenously compulsion (grooming) activated system. Hyperactive asymmetrical behaviors such as LCL2 and SCRL were observed in the WAR strain, either at baseline, microinjected with SAL or OT, similar to Adamec and Morgan [43] description of left hemisphere decreased anxiety after electrical amygdala kindling, in contrast with right hemisphere increased anxiety. This suggests an analogous expression of asymmetrical motor patterns in WARs, probably linked also to asymmetrical cognitive/emotional processing circuits.

PFC, ST, GP and TH are part of connected neural networks involved in alternating patterns and their dysfunctions are underlying OCD [22]. Furthermore, knock-out mice for the SAP90/PSD95-associated protein 3 (SAPAP3), postsynaptic scaffolding protein of excitatory synapses, densely expressed in ST and cortex, display compulsive grooming and anxiety [44]. [45], commented in [46], describe also excessive grooming in mutants of the homeodomain Hoxb8 and [47] showed that loss of neuron-specific transmembrane protein, SLIT and NTRK-like protein-5 (Slitrk5), induce excessive self-grooming and anxiety, with overactivation of orbitofrontal cortex. For a recent review on neurobiology of grooming see [48].

Therefore, although we do not have molecular studies looking for comorbidities and abnormalities in genes such as SAPAP3, Hoxb8 and Slitrk5 (associated to compulsive grooming) in WARs, [49] and [50] have shown that BTBR T+tf/J inbred mouse displays repetitive self- grooming and symptoms of autism, at the same time chronically seizing WARs display a decrease in the expression of hippocampal non-coding RNA BC1 [51], known repressor of the fragile X-associated protein [52,53], strongly associated to genetically-caused autism [54].

Moreover, [55] defined the so-called “*endogenous anti-convulsant system”*, detecting SNPr-associated muscimol-induced “*choreic-like”* behaviors, typical of Hungtingtons Chorea hyperkynetic activity, which blocked seizures. [56] also noticed with the anti-convulsant drug lamotrigine, a side-effect, called “*tourettism”*. Coincidently [57] described oro-facial automatisms and a cluster of grooming behaviors in Wistar rats resistant to acoustic stimulation. We therefore suggest that because of the tight inbreeding protocol (sister × brother; 60 generations) we observe nowadays genetic predisposition to seizures, and thanks to genetic linkage, WAR hypergrooming and networks genetically and concomitantly selected.

The correspondent Fos+ activation of both traditionally OCD-associated cortico-striatal-thalamic-cortical network, extended currently to a CeA-driven PVN-DG-SNPr, SNPc and PFC network, is in consonance with the description of subpopulations of CeA neurons that actively suppress cholinergic-mediated cortical arousal and exploratory behavior and simultaneously promote freezing responses, serving as a *switch* between active and passive fear [58]. Moreover, our data might be the experimental counterpart of the detection by positron emission tomography (PET) of activation in human CeA, in a provoked paradigm to elicited symptoms of OCD, particularly washing/cleaning associated to fear of contamination [59].

In the other hand, our study is a contribution to the need for detailed and rigorous behavioral studies, as mentioned by [60], in their recent claim “*Neuroscience Needs Behavior”*, in the context of the contemporary neuroscience literature, in which causality and molecular mechanisms are strongly proposed, even in the absence of behavioral phenomena definition and context, or when behavior and its rational are basically presented as secondary (appendix) material.

## 5. Conclusions

In conclusion, current data with behavioral (scores, flowcharts, graphs, Markov chains) and cellular (Fos) multivariate analysis, demonstrate that, in addition to epilepsy, WARs present hypergrooming, endogenously as in WAR-Basal, or induced by microinjection of OT-CeA, as a neuropsychiatric comorbidity depending on the activation of specific neural networks. Although the current behavioral alterations are driven from CeA, the consequent Fos+ neuronal networks are composed by PVN, PFC, ST, DG, SNPc and SNPr, important structures, part of well-known traditional OCD-associated cortico-striatal-thalamic-cortical circuits.

We are quite positive about the impact of these integrated behavioral, cellular and modeling approaches to experimental compulsion-epilepsy comorbidity, for neurology, neuropsychiatry and neuroscience.

## 6. Acknowledgments

We thank all the colleagues from the Neurophysiology and Neuroethology Laboratory. Special thanks to Prof. Edson Zangiacomi Martinez from the Center for Quantitative Methods (CEMEQ) at the Department of Social Medicine, Ribeiro Preto Medical School, University of So Paulo. Financial support: FAPESP-Cinapce (protocol 2005/56447-7) and Thematic Project (protocol 2007/50261-4), CAPES-PROEX (Neurology and Physiology FMRP), CNPq and FAEPA. NGC holds a CNPq Research Fellowship.

